# The Impact of *Arachis Hypogaea* Diet on Malaria Parasite Resistance and Haematology in Mice Infected with *Plasmodium berghei*

**DOI:** 10.64898/2026.04.03.716360

**Authors:** Obiageli A. Okeke, Goodness Daberechi Aniekwe, Onyebuchi Miracle Ndinyelum, Kingsley Chinemerem Mbelede, Cyril Ali Imakwu, Benedette Ifeoma Anyamene, Chukwuemeka Nwabunwanne Nwafe, Chizoba Favour Ndubuisi, Ifeyinwa Chinazom Ginikanwa, Nneka Lysa Kobune

## Abstract

This study evaluated the effects of different Arachis hypogaea dietary preparations on parasite load, haematological indices, and physiological responses in Plasmodium berghei-infected mice. Forty-five albino mice were randomly assigned to five groups: normal control, infected untreated control, roasted groundnut, boiled groundnut, and a combination of roasted and boiled groundnut diets. Data were analyzed using one-way ANOVA at p > 0.05. Infection resulted in a high parasite load in the untreated group, with no significant difference compared to the boiled and combined diet groups. However, the roasted groundnut group showed a reduction in parasite load and relatively higher chemosuppressive activity, although differences were not statistically significant. White blood cell counts increased significantly following infection, and dietary treatments did not restore normal levels. Similarly, red blood cell counts and packed cell volume were significantly reduced in infected mice. The roasted groundnut diet moderately improved PCV compared to other treatments but did not restore it to normal levels. Weight loss was most pronounced in untreated mice, while roasted groundnut intake showed slight mitigation. No significant effects on temperature regulation were observed. Overall, A. hypogaea diets did not significantly improve parasitemia or haematological parameters, indicating limited therapeutic value in malaria management.

**Importance:** This study is of significant importance due to its contribution to the ongoing search for accessible, affordable, and nutritionally based supportive interventions in malaria management. Malaria remains a major public health burden, particularly in sub-Saharan Africa, where increasing resistance of *Plasmodium* species to conventional antimalarial drugs continues to undermine control efforts. By investigating the effects of *Arachis hypogaea* (groundnut), a widely consumed and locally available food resource, this research explores a practical dietary approach that could complement existing malaria treatment strategies. In summary, this research is important because it bridges nutrition, parasitology, and public health, offering practical insights that could inform both scientific advancement and real-world malaria management strategies.

## Introduction

Malaria is a vector borne parasitic disease, transmitted by the female *Anopheles* mosquito carrying the *Pasmodium parasites*. Of all the species that can cause human malaria, *Plasmodium falciparum* and *Plasmodium vivax* are the most important (WHO, 2023) *P. falciparum* causes the most severe form of malaria and is commonly found in Africa, while P. vivax is the parasite of interest in many areas beyond sub-Saharan Africa (Nzekwe *et al*., 2024). It remains a major health challenge in the world due to its emergence and spread of parasite resistance to established antimalarial drugs (Wellems *et al*., 2001). Malaria disproportionately affects vulnerable populations, including children under five years of age and pregnant women, further exacerbating socio-economic disparities and hindering development efforts in endemic regions. The fight against malaria has been characterized by multifaceted approaches, ranging from vector control and case management to preventive measures and research into vaccines and novel interventions (Mitsakakis, *et al*., 2018).

The manifestations of malaria illness result from the infection of the erythrocytes and can reach host tissues through the circulatory system. The interactions between infected erythrocytes and host tissue are limited to locally specific tissue sites such as the brain, lung, liver, kidney, and intestines (Coban *et al*., 2018). A higher incidence of malaria occurs in the rainy season because more favourable conditions for mosquito breeding to occur (Nabatanzi *et al*., 2022). Severe malaria can lead to complications, such as anaemia, thrombocytopenia, leucocytosis, antioxidant reductions, increased lipid peroxidation, lactic acidosis, coma and death (Olayemi *et al*., 2012).

Groundnut or peanut (*Arachis hypogaea L*.) is one of the important edible oil seed Crop cultivated in world. Peanut plays an important role in the economy of several countries (Campos-Mondragón *et al*., 2009). Groundnut are high in oil content and compared with other major oil seeds, are relatively low in ash and carbohydrate (Tuberoso *et al*., 2007). Fatty acids, particularly oleic and linoleic, have a large bearing on the stability and nutritional quality of peanut oil. Groundnut plant known for its nutritional and health benefits with five main nutrients required by the body to maintain and repair the tissues namely energy, protein, phosphorus, thiamin, and niacin are found in good quantity in it (Singh and Kunwar, 2018). Groundnut is rich in vitamins and contains at least 13 different types of vitamin that include vitamin A, B, C and E, it is rich in 26 essential minerals including calcium, Iron, Zinc, boron.

All this plant nutrients are of great benefit to the body which brings about the study or the investigation of this Groundnut diets on malaria resistance and haematology in mice. Therefore, understanding of the host protective mechanisms against malaria and parasite-host interactions is essential, using dietary management strategies to control malaria.

## Materials and methods

### Procurement and Preparation of Food Materials

Groundnut and commercial pellet feed were purchased at the local Market in Awka, Anambra State, Nigeria.

### Preparation of Roasted Groundnut Flour

Groundnut was cleaned and sorted after which it was steamed and dried. It was then roasted above 100°C for 25 minutes in a gas oven, then dehulled, milled and sieved to obtain roasted groundnut flour.

### Preparation of Boiled Groundnut Flour

Groundnut was cleaned and sorted after which it was steamed also at the temperature 100°C in 45 minutes and dry for two weeks before been milled and sieved to obtain boiled groundnut flour.

### Procurement, Management of Experimental Animals and ethical approval

A total number of 45 Swiss albino mice with average weight of (21.91g) 10 weeks’ old were obtained from the Institute for Advanced Medical Research and Training (IAMRAT) college of medicine, University of Ibadan. Four mice (donor mice) were obtained from the Nigerian Institute of Medical Research, Lagos (NIMR). The mice were transported to the research area – the Zoology Laboratory in the Faculty of Biosciences, Nnamdi Azikiwe University, Awka in a transportation plastic cage measuring 40 × 20 × 20 cm to ensure adequate ventilation. The mice were acclimatized for a period of one week before the experiment began. They were housed in 15 cages in the Zoology Laboratory of the Faculty of Biosciences, Nnamdi Azikiwe University, Awka and fed with formulated food and water; morning and evening daily (Ndinyelum and Ufele-Obiesie, 2024). The vital chick mash was fed the animals; this feed is designed to provide young animals with a healthy growth and development. This is noted for its high concentration of protein, energy, vitamins and minerals as well as essential amino acids for protein synthesis and growth (Ndinyelum and Ufele-Obiesie, 2024). All animal experiments were conducted in compliance with NIH Guide for the Care and Use of Laboratory Animals and approval obtained from the University institutional animal care and use committee.

### Experimental Design

A cross sectional descriptive survey was conducted amongst adults in both urban and rural area. One hundred persons participated and the response was obtained using a questionnaire. Statistical analysis of the results gotten from the questionnaire was used to get the average amount of food material consumed and the frequency of consumption. From the results gotten the animal equivalent dose was calculated using the formula;

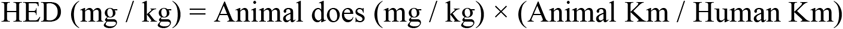

Where;

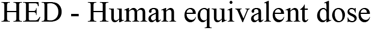

Km factor for each species is constant, the Km ratio is used to simplify calculations

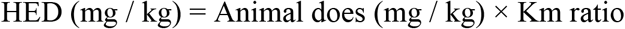

The Km ratio is gotten from the table below Source (Nair *et al*., 2016).

The experimental animals were randomly housed in 11 stainless steel metabolic cages laid out in a complete randomized design (CRD) of three treatments, replicated thrice with each replicate having three mice, and fed with commercial food (Vital Feed Broiler Starter, 18.00 ± 0.50g/100g crude protein, and 2106.00 kcal/kg metabolizable energy, Vital Feed, Grand Cereals Limited, Jos, Plateau State, Nigeria) and water ad libitum daily (Ndinyelum and Ufele-Obiesie, 2024).

The experimental animals which were of homogenous sizes were randomly stocked into three cages (24 by 24 cm) containing three mice each and labelled as follows; Group 1: Normal Control (Uninfected, Untreated); Group 2: Negative Control (Infected, Untreated); Group 3: were fed with 14g of roasted groundnut and 22g of commercial feed. Prior to and after infection; Group 4: were fed with 14g of boiled groundnut and 22g of commercial feed. Prior to and after infection; Group 5: were fed with 7g of roasted groundnut and 7g of boiled groundnut and 22g of commercial feed. Prior to and after infection.

### Parasite inoculation

The infected erythrocytes for each test were collected from the infected donor mice with rising parasitemia of 20 to 30%. The mice were sacrificed through cardiac puncture, and blood was collected in a heparinized tub with an anticoagulant (0.5% trisodium citrate). The blood was then diluted with normal saline (0.9%) in proportion of 1:4. Each mouse in every group was then inoculated with 0.2 mL of blood containing about 1.25×10^6^ *P. berghei* infected erythrocytes two weeks after feeding them with experimental feeds through intra peritoneal route.

### Determination of Parasite load

Parasitemia was measured by collection of blood sample through tail bleeding for the preparation of thick and thin film smear and stained with Giemsa-stain. Parasitemia was calculated based on the number of infected erythrocytes in counted erythrocytes observed. The formula for parasitemia is as follows:

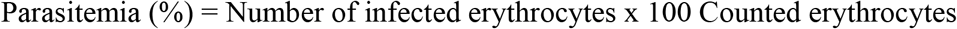

The percentage of inhibition was calculated using the following formula:

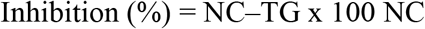

NC: mean of parasitemia in negative control

TG: mean of parasitemia in treated-group (G1, G2, G3).

### Body weight gain

The weights of the mice were measured and recorded using a sensitive weighing scale, after which the mice were randomly placed into different sections of the cage within the range of the body weight gain per mice for the different treatment groups (Duwa *et al*., 2020). Weight gain will be calculated as the difference between two successive weekly body weights as follows:

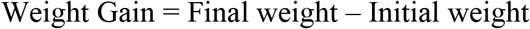

### Temperature

The temperature of the mice was measured and recorded using a thermometer, after which the mice were randomly placed into different sections of the cage, this was done before and after acclimatization.

### Determination of Packed Cell Volume (**PCV**)

The packed cell volume was determined making use of the micro haematocrit centrifuge method as described by (Farooq *et al*., 2023).

### White Blood Cell (**WBC**)

The WBCs were counted using the principle of calibrated capillary tube for blood sampling with Haemocytometer (Ndinyelum and Ufele-Obiesie, 2024).

Number of WBC in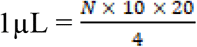. Source: Verso (1964).

Where N; is the number of cells counted,

10; is the depth of the counting chamber, 20; is the dilution ratio – 1:20, and

4; is the squares counted.

This is summarized into N× 50; which gives the value of WBC.

### Red Blood Cell (**RBC**)

The RBC count was determined using the principle of a calibrated tube for blood sampling with haemocytometer method (Ufele-Obiesie and Ndinyelum, 2025).

Number of RBC in 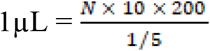. Source: Verso (1964).

Where N; is the number of cells counted,

10; is the depth of the counting chamber, 200; is the dilution ratio – 1:200, and

5; is the squares counted.

This is summarized into N× 10000; which gives the value of RBC.

### Statistical analysis

The data of haematological indices, weight, temperature and parasitemia of mice were subjected to mean, standard deviation and One-Way Analysis of Variance (ANOVA) was carried out using SPSS (25) (IBM SPSS, 2017). The Tukey’s Honestly Significant Difference was used to separate mean significant differences between treatments at a 5% significant level (Okeke and Mogbo, 2013). Results expressed as Mean+SD; Mean values with different alphabets as superscripts are significantly different (p<0.05).

## RESULITS

### Impact of *A. Hypogea* on malaria parasite resistance

The mean parasite load in the infected and untreated mice was 6.77 ± 0.06% (range: 6.70% to 6.80%). Treatment with a combination of roasted and boiled *A. hypogea* resulted in a mean parasite load of 6.40 ± 2.60% (range: 4.80% to 9.40%), which did not significantly affect the parasite load. Similarly, treatment with single diets of roasted *A. Hypogea* (mean: 1.47 ± 0.31%, range: 1.20% to 1.80%) and boiled *A. hypogea* (mean: 6.00 ± 3.70%, range: 3.80% to 10.80%) also did not significantly impact the parasite load in the infected mice (Figure 1).

**Figure 1.**
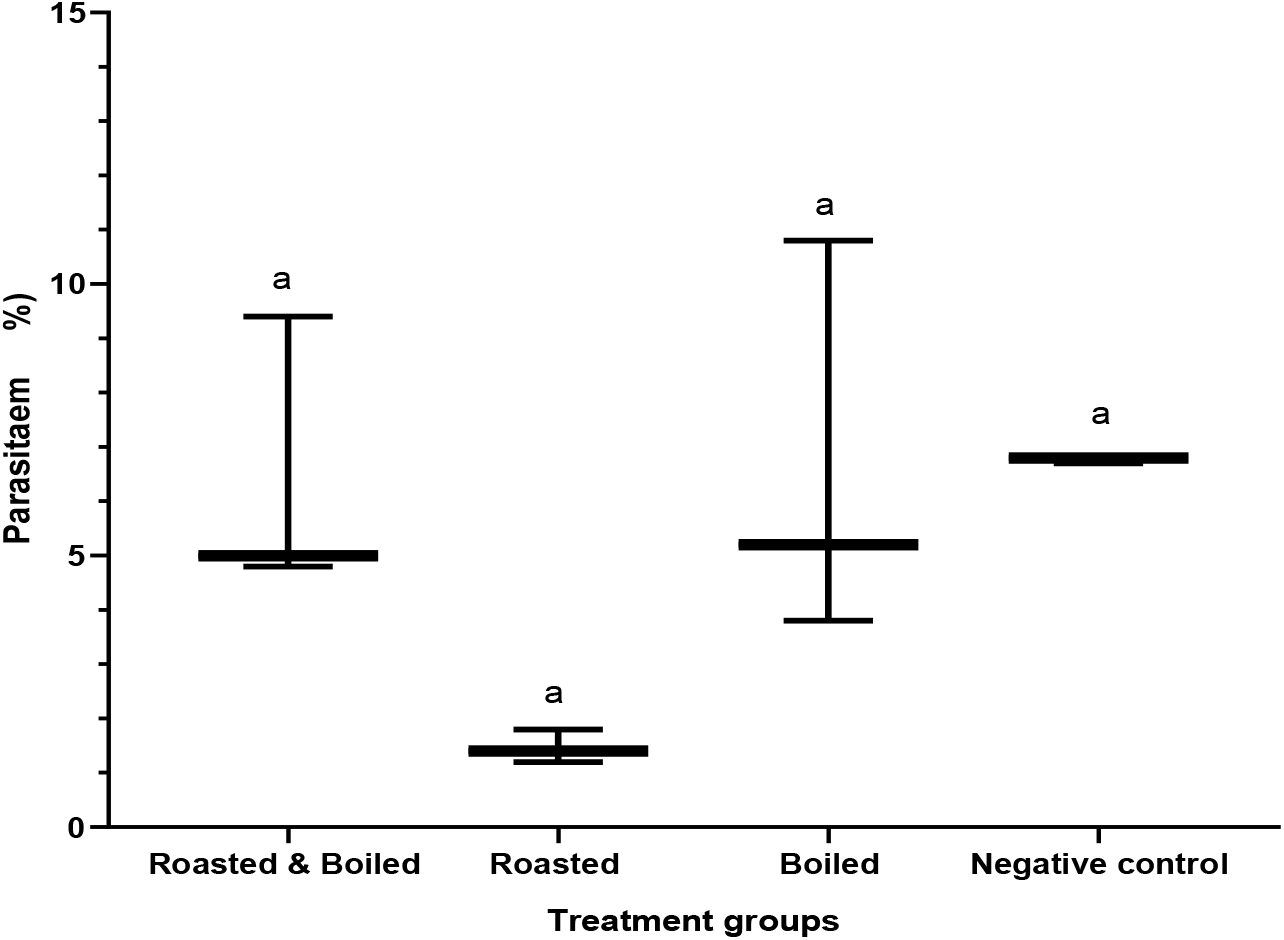
Impact of *A. hypogea* on parasite load of *P. berghei*-infected mice Chemosuppressive activity of *A. hypogea* on parasite load of *P. berghei*-infected mice.

The diet consisting of roasted *Arachis hypogea* demonstrated the strongest chemosuppressive activity against *Plasmodium berghei* infection, with a mean suppression rate of 78.43% (ranging from 72.53% to 82.35%). However, this activity did not significantly differ from the other treatment groups fed *A. hypogea*. In the group that received a combination of roasted and boiled *A. hypogea*, the mean chemosuppressive activity was only 5.88% (with a range from -38.24% to 27.41%), while the group fed only boiled *A. hypogea* showed an even lower mean activity of 2.9% (ranging from -58.82% to 44.12%). Overall, there was no significant difference in the chemosuppressive activity among all the treatment groups (Figure 2).

**Figure 2.**
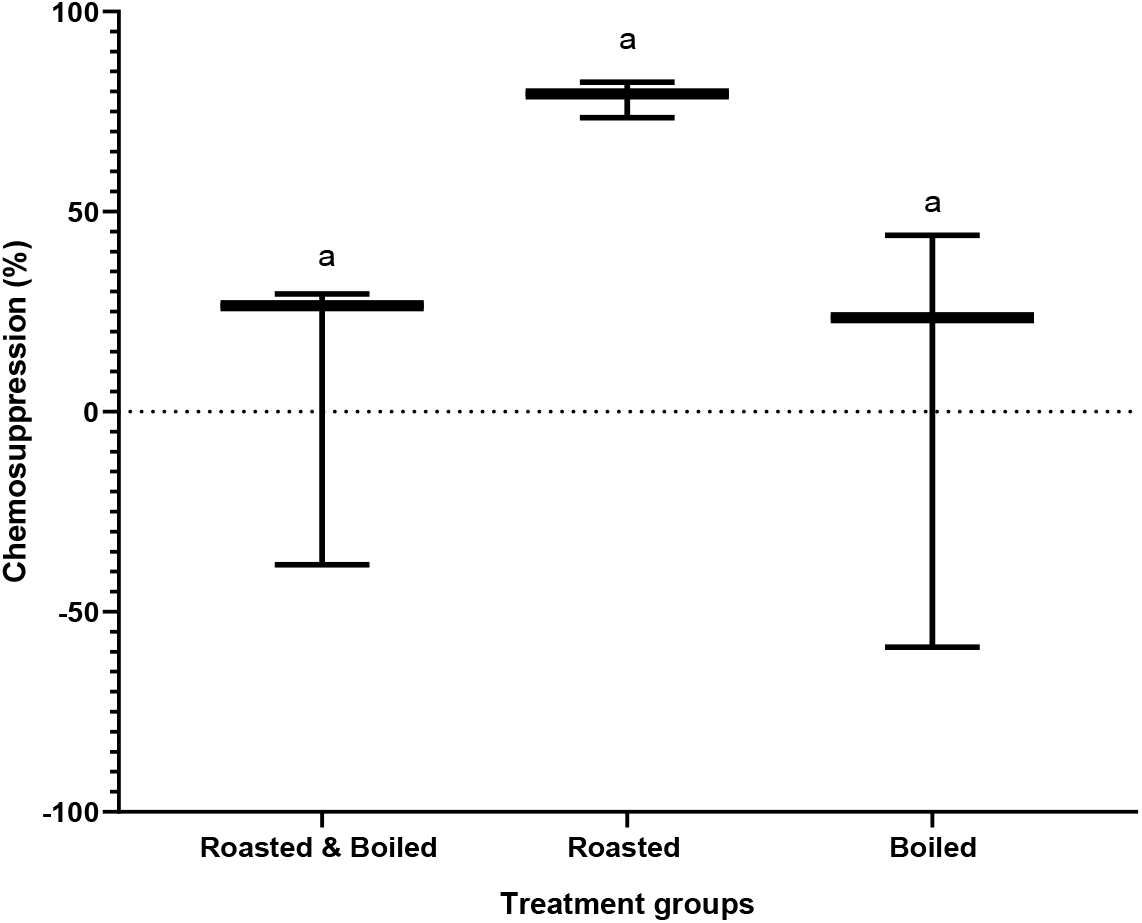
Chemosuppressive activity of *A. hypogea* diets on *P. berghei*-infected mice Impact of *A. hypogea* diets on WBC of *P. berghei-*infected mice.

Infection with the parasite resulted in a significant increase (p < 0.05) in white blood cell (WBC) counts in the negative control group compared to the normal control group. However, treatment with the various groundnut diets is not significantly different (p > 0.05) with WBC levels to a degree comparable to each other (Figure 3).

**Figure 3.**
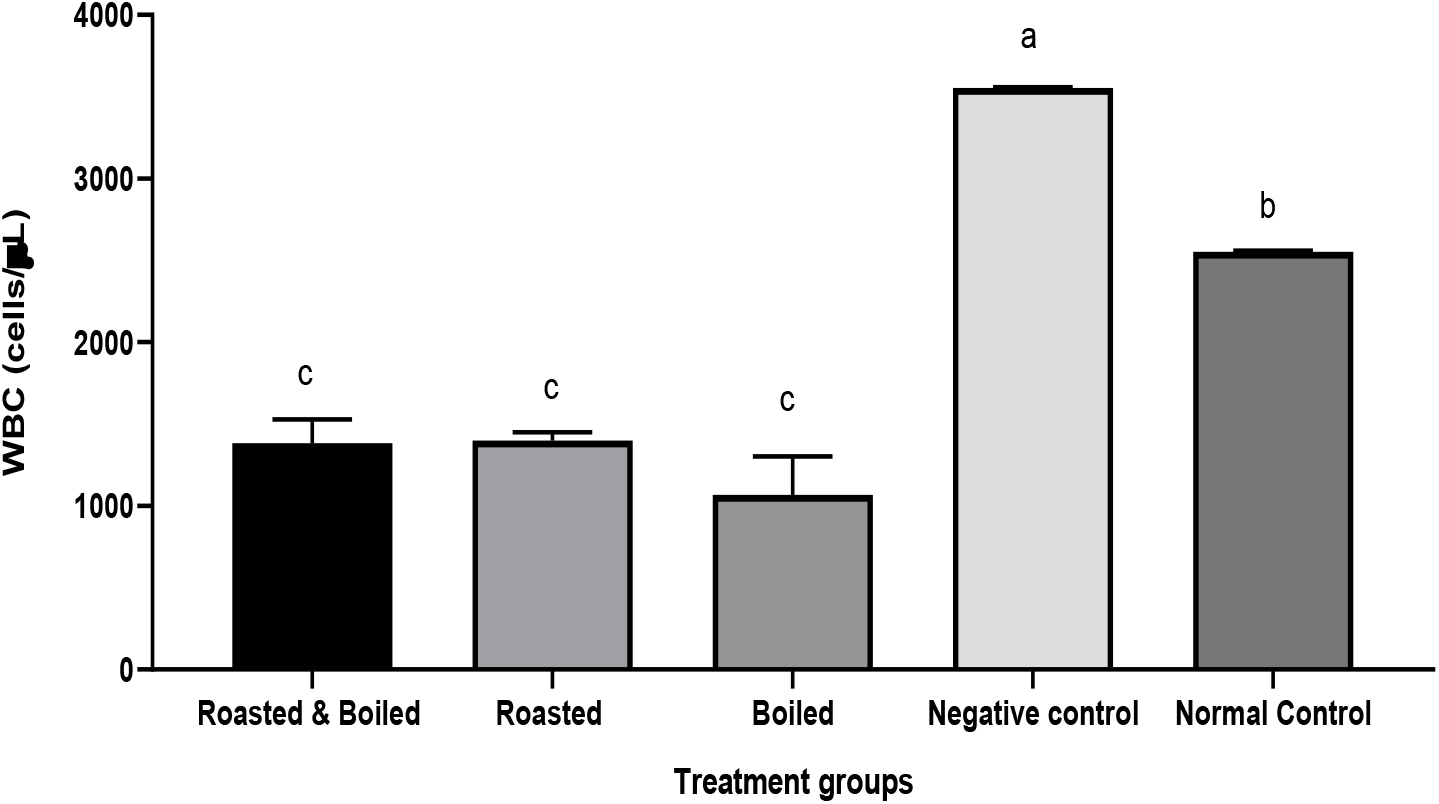
Impact of *A. hypogea* diets on WBC of *P. berghei*-infected mice Impact of *A. hypogea* diets on RBC of *P. berghei*-infected mice.

The red blood cell (RBC) count in normal mice was significantly higher than in the other treatment groups. Infection with the parasite caused a notable decrease (p<0.05) in RBC levels compared to the normal group. While treatment with roasted groundnuts aimed to increase the RBC in the mice, the results were not comparable (p<0.05) to the normal group, and the RBC remained below (p<0.05) that of the negative control. Additionally, other groups fed different groundnut diets did not show any improvement (p<0.05) in RBC levels (Figure 4).

**Figure 4.**
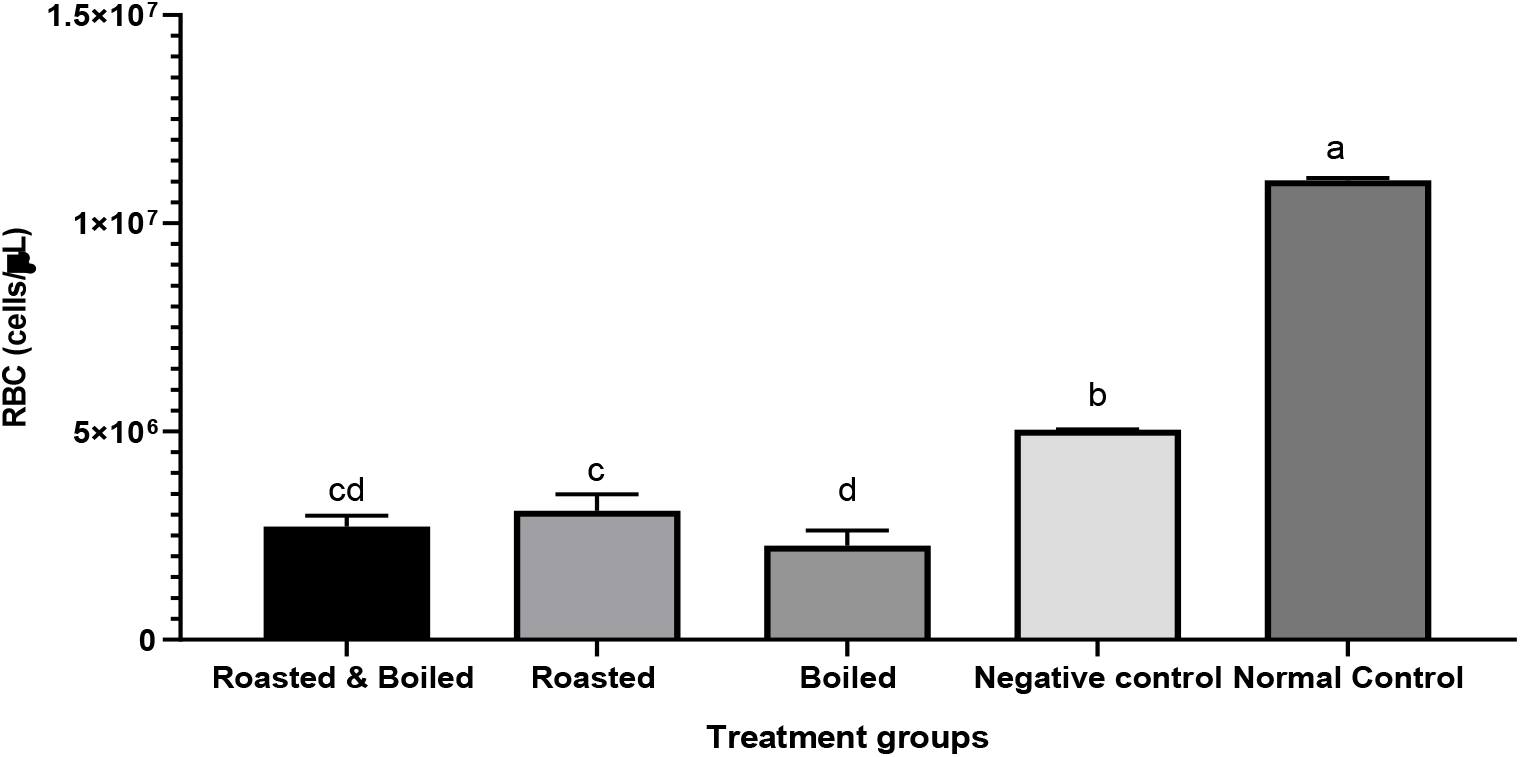
Impact of *A. hypogea* diets on RBC of *P*.*berghei*-infected mice Impact of *A. hypogea* diets on PCV of *P. berghei*-infected mice.

The packed cell volume (PCV) of the normal group was significantly higher (p < 0.05) compared to other treatment groups. Infection with the parasite caused a significant decrease (p < 0.05) in parasite load. The PCV of the group fed roasted groundnuts was significantly higher (p < 0.05) than those of the groups fed a combination of roasted and boiled groundnuts and boiled groundnuts only. However, feeding infected mice different groundnut diets did not improve (p > 0.05) the PCV to levels above the negative control or bring it in line with the normal group (Figure 5).

**Figure 5.**
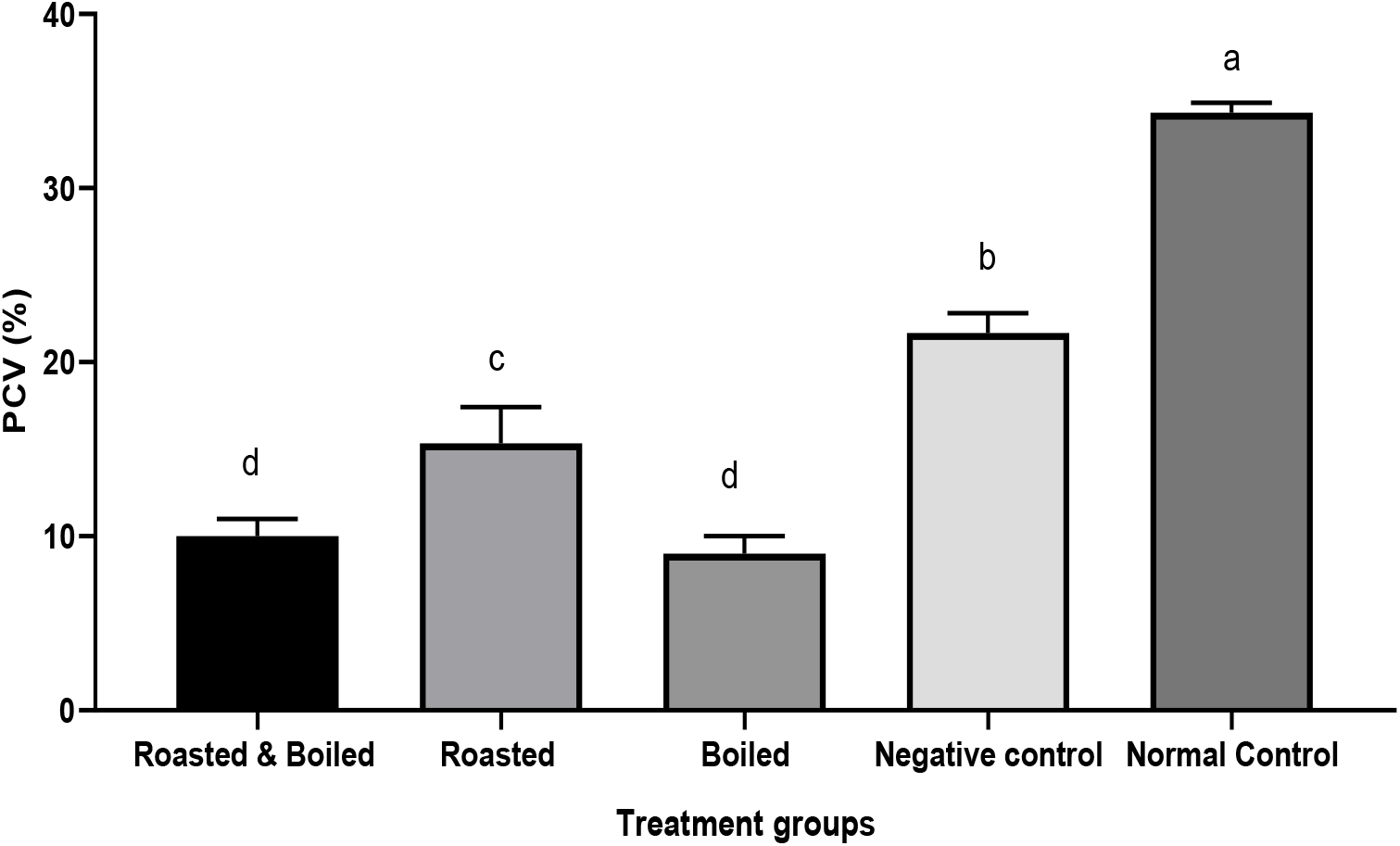
Impact of *A. hypogea* diets on PCV of *P. berghei*-infected mice Impact of *A. hypogea* diets on Body Weight and Temperature of *P. berghei*-infected mice.

The negative control group experienced significantly greater weight loss (p<0.05). Mice treated with roasted *A. hypogea* showed an increase in weight that was significantly higher than that of the negative control group; however, this increase was not substantial enough to differ significantly from the other *A. hypogea*-fed groups. In the groups of mice that were fed a combination of roasted and boiled *A. hypogea*, as well as those that received boiled *A. hypogea* only, weight loss was observed (Table 2). Among the groups fed *A. hypogea*, only the group that consumed the roasted form of the diet demonstrated a temperature change that was comparable to normal levels at the end of the study (p>0.05) (Table 2).

**Table 1.**
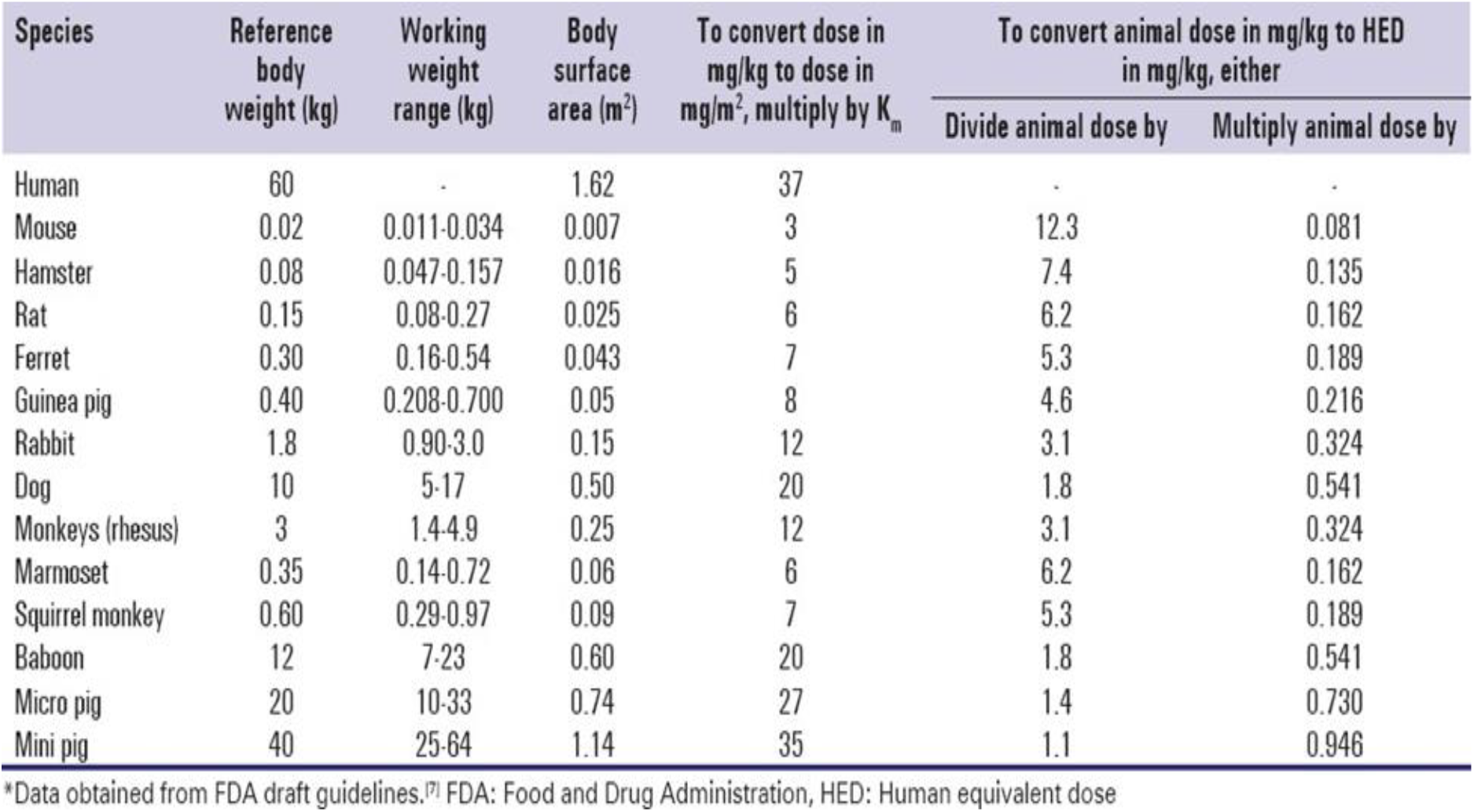
HED conversion table.

**Table 2.** Impact of *A. hypogea* diets on body weight and temperature of *P. berghei*-infected mice.

| Groups | Weight of animals (g) | | | Temperature of animals ($^{\circ}\text{C}$) | | |
| --- | --- | --- | --- | --- | --- | --- |
|  | Initial | Final | Change in weight | Initial | Final | Change in Temp. |
| Roasted<br>& Boiled | 27.77±1.45<sup>cd</sup> | 26.97±0.55<sup>c</sup> | -0.80±0.90<sup>b</sup> | 37.00±0.90<sup>a</sup> | 38.30±0.50<sup>a</sup> | 1.30±0.40<sup>b</sup> |
| Roasted | 30.47±1.85<sup>bc</sup> | 30.77±1.65<sup>b</sup> | 0.30±0.20<sup>b</sup> | 37.07±0.85<sup>ab</sup> | 37.80±0.60<sup>ab</sup> | 0.70±1.45<sup>ab</sup> |
| Boiled | 26.47±0.06<sup>d</sup> | 25.00±1.30<sup>c</sup> | -1.47±1.35<sup>b</sup> | 37.50±0.10<sup>b</sup> | 35.87±1.85<sup>b</sup> | -1.63±1.75<sup>b</sup> |
| Negative<br>Control | 37.63±0.06<sup>a</sup> | 30.33±0.06<sup>b</sup> | -7.30±0.10<sup>c</sup> | 37.43±0.06<sup>a</sup> | 36.83±0.06<sup>a</sup> | -0.60±0.10<sup>ab</sup> |
| Normal<br>Control | 31.77±0.06<sup>b</sup> | 35.37±0.06<sup>a</sup> | 3.60±0.00<sup>a</sup> | 38.93±0.06<sup>a</sup> | 39.07±0.12<sup>a</sup> | 0.13±0.06<sup>ab</sup> |

## Discussion

The present study evaluated the effects of differently processed *Arachis hypogaea* diets on parasitemia, haematological parameters, and physiological indices in *Plasmodium berghei*-infected mice. The infected untreated group exhibited high parasite load with minimal variability, indicating consistent infection across subjects. Although the combined roasted and boiled diet showed a slight reduction in mean parasite load, the wide variability suggests no significant impact on parasite burden, consistent with reports that high-fat diets may modulate immune responses rather than directly reduce parasitemia (Soares *et al*., 2023; Cimperman *et al*., 2023).

Roasted *Arachis hypogaea* diet showed the highest mean chemosuppressive activity against *Plasmodium berghei* infection indicating a substantial reduction in parasite levels on average. The range is quite narrow, showing consistent suppression effectiveness within this group. Combination of roasted and boiled *A. hypogea* shows much suppression lower with a wide range. Boiled *A. hypogea* exhibited the lowest mean chemosuppressive activity with an even wider and more variable range. The consistent and high suppression achieved by the roasted groundnut compares favourably with boiling groundnut suggesting reliable anti-plasmodial properties. Similar findings were reported by (Chandel and Bagai, 2010), where ethanolic leaf extracts of other plant treatment; *Ajuga bracteosa* showed dose-dependent chemosuppression of *Plasmodium berghei* in mice with up to 77.7% suppression at higher doses and prolonged survival, indicating substantial parasite reduction similar to roasted *Arachis hypogaea*’s effect.

Infection caused a significant increase in white blood cell (WBC) counts in the negative control group compared to the normal control group. Similar finding on the effects of oil diet reports that several studies showing that infections cause a significant increase in white blood cell (WBC) counts compared to control groups (Okeke *et al* 2026; Polis, *et al*., 2024; Xu, *et al*., 2013). Antibodies which is produced by the WBC can act on restricting the growth of blood stage parasites, bringing about their increase and the progression of clinical symptoms for several mechanisms, like obstructing erythrocyte invasion (Innocent, *et al*., 2024). Groundnut diets did not improve immune normalization as measured by WBC counts, implying limited effect on controlling or modulating systemic inflammation or immune activation triggered by the infection. This aligns with (Innocent *et al*., 2024), that indicates groundnut diets did not improve immune normalization as measured by white blood cell (WBC) counts, suggesting limited effects on controlling systemic inflammation or immune activation triggered by infection.

Normal (uninfected) mice have the highest RBC counts, serving as baseline healthy control. *Plasmodium berghei* infection causes reduction in RBC counts which can bring about anaemia/haemolysis due to parasitic infection (Ekaidem and Henry, 2016), this aligns with established knowledge that *P. berghei* infection causes anemia in mice by reducing RBC counts through a combination of haemolysis of infected RBCs and suppression of RBC production due to inflammatory responses. In this study normal mice serve as the healthy baseline animals with highest RBC counts before infection-induced declines occurred (Lakkavara *et al*., 2020; Junaid *et al*., 2017).The infection clearly induced a statistically significant decrease in RBC levels compared to normal mice. This is expected as malaria parasites invade and destroy RBCs, leading to anaemia (Dalimot *et al*., 2022).

Impact of Roasted Groundnut Diet aims to increase PCV, thereby prevents or reduce anaemia but the PCV remained below that of normal mice which indicates that the roasted groundnut diet was unable to fully restore RBC volume or count to healthy levels during infection. This is similar to prior study by (Innocent, *et al*., 2024), which also reported that the roasted groundnut diet effects on PCV during infection is beneficial but insufficient to restore PCV volume fully to healthy baseline levels (Isamoh *et al*., 2013). The result from this study showed that the normal group has the highest PCV. Among infected groups, the group fed roasted groundnuts showed a higher PCV (p < 0.05) than the groups fed combination roasted and boiled groundnuts or boiled groundnuts only. This shows that roasted groundnuts better preserved PCV relative to the other dietary treatments. In support to (Innocent *et al*., 2024), shows that the PCV percentage for fresh groundnut group was the highest, followed by fried, controlled and boiled groundnut groups, this shows that roasted groundnuts may offer some moderate benefit in mitigating PCV reduction caused by infection, the improvement is insufficient to fully reverse or prevent anaemia associated with the parasite infection.

The negative control group (infected and untreated) experienced greater weight loss compared to at least some other groups. This align with Schwarz, *et al*. (2011). This indicates that infection or lack of treatment adversely affected their weight. Mice treated with roasted *A. hypogea* showed a significant increase in weight relative to the negative control, consistent with Soarer *et al*. (2023) report, suggesting a protective or restorative effect of this diet in terms of weight maintenance or gain. The groups fed combination (roasted and boiled) and boiled only *A. hypogaea* diets experienced weight loss, indicating less beneficial on weight compared to the roasted-only diet.

Only the group fed the roasted *A. hypogaea* diet showed a body temperature change that was comparable to normal (non-infected or healthy) levels by the end of the study. There is no direct or specific research found that confirms or contradicts the observation that only the group fed a roasted *A. hypogaea* (peanut) diet showed body temperature changes comparable to normal (non-infected or healthy) individuals. Existing studies mostly focus on the nutritional and sensory effects of roasting peanuts, demonstrating that roasting at moderate temperatures preserves nutrients and enhances flavor without any negative impacts. While roasting peanuts improves nutritional quality and flavor, the specific effect of roasted peanut diets on body temperature remains unverified and requires further targeted research (Zhang *et al*., 2024).

## Conclusion

This study demonstrates that roasted *Arachis hypogaea* diet exhibits moderate antiplasmodial and supportive physiological effects in *Plasmodium berghei*-infected mice, outperforming boiled and combined preparations. However, none of the dietary interventions effectively restored parasitemia or haematological parameters to normal levels. These findings suggest that while roasted groundnut may provide supportive benefits, it is insufficient as a standalone therapy for malaria. Further studies should explore its potential synergistic effects with standard antimalarial drugs and investigate the mechanisms underlying its bioactivity.

## Acknowledgement

The researchers are thankful to the laboratory technologist and other lecturers in the department who contributed to this research.

